# ^18^F-FDG-PET Hyperactivity in Alzheimer’s Disease Cerebellum and Primary Olfactory Cortex

**DOI:** 10.1101/2020.06.05.136838

**Authors:** Mark D. Meadowcroft, Carson J. Purnell, Jian-Li Wang, Prasanna Karunanayaka, Qing X. Yang, The Alzheimer’s Disease Neuroimaging Initiative

## Abstract

Cerebellar involvement in Alzheimer’s disease (AD) has not been studied to the extent that cortical neuropathological changes have. Historical and recent histopathological literature demonstrate cerebellar AD pathology while functional investigation has demonstrated disrupted intrinsic cortical – cerebellar connectivity in AD. Additionally, olfactory deficits occur early in AD, prior to the onset of clinical symptoms. The neurological basis for the involvement of the cerebellum and olfactory system in the disease course remain unclear. ^18^F-fludeoxyglucose (FDG) positron emission tomography (PET) data from the Alzheimer’s Disease Neuroimaging Initiative (ADNI) were analyzed to characterize metabolism in the cerebellum and olfactory region of AD, mild-cognitive impaired (MCI), and age-matched cognitively normal (CN) controls. In contrast to known parietal and temporal lobe FDG *hypo-metabolism* within the default mode network in AD, a significant FDG *hyper-metabolism* was found in the cerebellum and olfactory cortical regions (including the piriform cortex, olfactory tubercle, anterior olfactory nucleus, and nucleus accumbens shell). The increase in cerebellum glucose utilization was shown also in late- verses early-MCI patients. The cerebellar and olfactory regions both contain inhibitory distal and inter-neuronal connections that are vulnerable to disruption in AD. The hyper-metabolism in the cerebellum and olfactory structures may reflect disruption of local and system-wide inhibitory networks due to AD neurodegeneration, suggesting a hypothetical mechanism for susceptibility of the olfactory system to early AD pathology.

## 1. Introduction

Fludeoxyglucose positron emission tomography (FDG-PET) quantification of cerebral glucose metabolism has been established as an *in-vivo* imaging marker for neurodegeneration related to Alzheimer’s disease (AD) [1]. FDG-PET is routinely used to differentiate AD from frontal temporal (FTD) and other dementias based on the distribution or pattern of regional brain cerebral glucose metabolic rate (CMRgl) with a high-degree diagnostic accuracy, sensitivity, and specificity [2, 3]. In AD patients images of FDG-PET commonly present as hypo-metabolic in the parieto-temporal (inferior parietal lobule), posterior cingulate cortex (PCC), and medial temporal lobes (MTL) [4]. This hypo-FDG activity in the MTL occurs early in the disease process and is evident in amnestic mild cognitively impaired (MCI) patients [5].

The hypo-FDG activity observed in AD has been attributed to a reduction in glucose utilization (hypo-metabolism) due to a loss of functional activity and is an index of neuronal/synaptic dysfunction, rather than simply an artifact of reduced neural cell count (atrophy) [6]. Brain atrophy is preceded and exceeded by hypo-metabolism in presymptomatic and prodromal AD and MCI in most regions, suggesting a multifactorial degenerative process [7, 8]. Hypo-metabolism is frequently observed in AD brain regions which are positive for amyloid-beta (Aβ) imaging, such as the parieto-temporal association cortices, posterior cingulate cortex, and the precuneus [9]. These regions are collectively part of a resting state brain network identified as the default mode network (DMN) [10]. Numerous studies have shown that DMN functional connectivity is reduced in AD. Since FDG-PET measures of glucose consumption and resting-state fMRI (rs-fMRI) metrics represent base-line neural activity, the hypo-metabolism and diminishing DMN functional connectivity in AD demonstrate a coupling between glucose utilization and transmission of neural information; brain glucose hypo-metabolism is linked to a reduced communication between brain regions impacted by underlying pathological AD processes [11].

Histopathological literature support cerebellar involvement in AD [12, 13]. Similar work has shown that there are selective vulnerabilities of the cerebellum to volumetric atrophy [14], amyloid deposition, late-stage gliosis, and neuronal loss in AD [12]. However, the involvement of the cerebellum has largely been overshadowed by study of cortical neuropathological changes in AD. While the cerebellum has long been regarded as a regulator of fine motor activity and learning; recent study has shifted our understanding of the cerebellum also towards a role in cognitive and emotional modulation [15]. The cerebellum is parcellated into discrete regions with functional connectivity to known cerebral brain networks, such as the default mode and salience networks [16]. Disrupted cerebellar-cerebral functional connectivity is observed in AD [17]. Somatosensory aspects of the cerebellum occupy a fraction of the whole, with the majority of the cerebellum networks outlined as having relevance in cognitive and emotional control [18].

From behavioral studies, prevalent olfactory deficits have long been observed in AD patients [19]. In addition to age-related loss of smell (presbyosmia) [20], olfactory dysfunction is found to precede the clinical manifestation of cognitive symptoms [21] and with increased incidence in individuals at higher risk of developing AD, such as primary relatives [22], APOe4 carriers [23], and MCI [24]. Pathologically, significantly higher amyloid plaque and neurofibrillary tangle load are observed in the olfactory bulb and olfactory cortex of Braak stage-I and II AD brains; suggesting a plausible hypothesis that AD-related pathology may begin in the primary olfactory structures such as the trans-entorhinal cortex and anterior olfactory nucleus and spread to MTL regions as the diseases progress [25].

Our prior olfactory fMRI studies demonstrated significant reductions in brain activity and connectivity in the primary olfactory cortex (POC) and related structures in AD and MCI subjects [21, 26, 27]. Thus, the olfactory deficits and neurodegeneration of the central olfactory system should play a central role in AD initiation and progression. However, as an AD neurodegeneration marker, the status of FDG-PET in the early AD pathology site of the olfactory structures has not been carefully examined. Similarly, cerebellum has been implicated in AD but has not been evaluated on a functional level. In order to understand the relationship between cerebellar changes, olfactory deficits, and neurodegeneration in AD, we evaluated resting-state FDG-PET metabolic activity in cerebellar and olfactory structures utilizing the Alzheimer’s Disease Neuroimaging Initiative (ADNI) data. Our analyses revealed remarkable characteristics in FDG-PET metabolism that provide a unique perspective for AD neurodegeneration.

## 2. Materials & Methods

### Patient selection

Data used in the preparation of this manuscript were obtained from the ADNI database (adni.loni.usc.edu), which includes the ADNI 1, GO, and 2 studies. The ADNI was launched in 2003 as a public-private partnership with the primary goal to test whether serial magnetic resonance imaging, positron emission tomography, other biological markers, and clinical and neuropsychological assessment can be combined to measure the progression of mild cognitive impairment and early Alzheimer’s disease. For up-to-date information, see www.adni-info.org.

A group of 245 ADNI subjects were utilized in this study as their data contained FDG positron emission scans and demographics with genotyping information. Patients were stratified based on their cognitive status: cognitively normal (CN) n=80, MCI n=149, and AD n=16 (Table 1). The MCI subject group was further sub-divided into early-(N=115) and late-MCI (N=34) status based on education adjusted ranges on the Logical Memory II subscale from the Wechsler Memory Scale (outlined in the ADNI-2 Procedures Manual [28]).

**Table 1.** Demographics, and Baseline Cognitive/Functional Test Measures

Demographics, and Baseline Cognitive/Functional Test Measures
| Demographic and Baseline Cognitive Tests | Alzheimer's Disease | Mild Cognitive Impaired |  | Cognitively Normal | ANOVA P Value |
| --- | --- | --- | --- | --- | --- |
|  |  | Late | Early |  |  |
| Gender | 5F, 11M | 16F, 18M | 57F, 58M | 42F, 38M | $\chi^2 < 0.001^*$ |
| Age (y) | 72.81 ± 2.76 | 74.06 ± 1.32 | 71.32 ± 0.66 | 75.67 ± 0.60 | < 0.001† |
| Education (y) | 16.50 ± 0.57 | 16.23 ± 0.39 | 15.72 ± 0.25 | 16.61 ± 0.29 | = 0.058‡ |
| CDR | 5.09 ± 0.29 | 1.87 ± 0.20 | 1.20 ± 0.07 | 0.03 ± 0.02 | < 0.001Δ |
| MSME | 22.88 ± 0.49 | 27.88 ± 0.26 | 28.39 ± 0.14 | 28.95 ± 0.14 | < 0.001α |
| MOCA | 16.13 ± 1.44 | 21.32 ± 0.51 | 23.49 ± 0.38 | 25.42.7 ± 0.42 | < 0.001Δ |
| ADAS-11 | 20.25 ± 1.93 | 12.38 ± 0.83 | 8.28 ± 0.32 | 5.92 ± 0.29 | < 0.001Δ |
| ADAS-13 | 30.81 ± 2.40 | 19.59 ± 1.15 | 13.06 ± 0.50 | 9.31 ± 0.43 | < 0.001Δ |
| RAVLT Immediate | 21.69 ± 2.24 | 31.62 ± 1.47 | 38.13 ± 0.91 | 44.90 ± 1.05 | < 0.001Δ |
| RAVLT Learning | 2.31 ± 0.49 | 3.56 ± 0.39 | 5.33 ± 0.22 | 5.96 ± 0.28 | < 0.001β |
| RAVLT % Forgetting | 78.19 ± 7.46 | 70.80 ± 4.12 | 46.31 ± 2.69 | 36.95 ± 2.62 | < 0.001Δ |
| Functional Assessment Questionnaire | 14.63 ± 1.47 | 4.03 ± 0.77 | 1.77 ± 0.29 | 0.10 ± 0.04 | < 0.001Δ |
| Everyday Cognition-Patient | 2.14 ± 0.18 | 1.82 ± 0.09 | 1.83 ± 0.51 | 1.32 ± 0.03 | < 0.001γ |
| Everyday Cognition-Study Partner | 3.02 ± 0.09 | 2.04 ± 0.11 | 1.64 ± 0.05 | 1.11 ± 0.03 | < 0.001Δ |

Only patients from the ADNI cohort with $^{18}\text{F}$ FDG-PET scan and genetic SNP data are included.
CDR = Clinical Dementia Rating, MMSE = Mini Mental State Examination, MOCA = Montreal Cognitive Assessment, ADAS-11 = Alzheimer's Disease Assessment Scale-Cognitive 11 Question Subscale, ADAS-13 = Alzheimer's Disease Assessment Scale-Cognitive 13 Question Subscale. RAVLT = Rey Auditory Verbal Learning Test
\* = AD patient gender distribution was different than CN and MCI, $\chi^2 < 0.001$ .
† = Gender was used as a factor in this analysis; a gender bias was determined with males overall having being older at baseline (74.24 ± 0.63 vs. 72.15 ± 0.67, $p = 0.023$ . CN > MCI, $p < 0.001$ .
‡ = Gender was used as a factor in this analysis; a gender bias was determined with males overall having more years of education (16.63 ± 0.22 vs 15.62 ± 0.24, $p = 0.002$ ).
$\Delta$ = age, gender, and education corrected. All between group comparisons $p < 0.001$ .
$\alpha$ = age, gender, and education corrected. Least significant between group comparison $p = 0.007$ .
$\beta$ = age, gender, and education corrected. Least significant between group comparison $p = 0.007$ .
$\gamma$ = age, gender, and education corrected. Least significant between group comparison $p = 0.001$ .

### Positron emission tomography (PET) protocol

FDG-PET data were obtained from the ADNI study that were previously acquired using specified parameters and CMRgl Patlak preprocessing [29]; all data can be found and downloaded from the LONI webserver. In short, subjects imaged in the morning were fasted overnight; those scanned in the afternoon were fasted at least four hours prior to the imaging session. Patients were staged in a standardized environment and asked to remain still and keep awake with eyes open looking straight ahead. 185 MBq (5 mCi +/− 10%) of [^18^F]-FDG was drawn, assayed with a dose calibrator, injected, flushed with 10 ml of saline, and time/dose recorded to the nearest minute. PET scanner acquisition parameters were as followed, scan start time post-injection = 30 min, dynamic 30 minute scan with six 5-minute frames, grid = 128×128, FOV = 256×256mm (2mm voxel size), and slice thickness = 3.27 mm.

### PET data processing

Individual FDG datasets were processed with AFNI (Analysis of Functional NeuroImages) [30]. Each 30-minute FDG time series was motion-corrected and averaged across all frames. Each patients FDG volume was partial volume corrected with a Van-Cittert deconvolution technique to improve quantitative accuracy and recover PET signal in cortical grey matter [31]. Each average was spatially normalized to a Montreal Neurological Institute (MNI) tracer template, resampled to 2 mm-isotropic, and the resulting image was used to standardize the uptake per subject. Each individual dataset was globally divided by the average voxel value in a whole-brain and cerebellar white matter mask to generate an SUVR image. Whole brain sub cortical white matter was selected as a reference region due to improved longitudinal cohort signal preservation [32, 33]. Images were smoothed with a 6 mm full-width half maximum (FWHM) Gaussian filter before parametric map analyses in SPM12. The SPM toolbox MarsBaR (MARSeille Boîte À Région d’Intérêt) [34] was used to output average values for ROI analysis from processed images. Regions of interest were input from the Automated Anatomical Labeling 2 (AAL2) [35] atlas normalized in MNI template brain space, consisting of 120 individual gray matter ROIs.

### Statistical design

Group-wise voxel based analyses of FDG-SUVR parametric maps was done in SPM12 using age and gender as covariates with cognitive diagnostic (AD, MCI, CN) as the variables of interest. MarsBaR outputs and demographic information for each group were compared with a MANCOVA (Multivariate Analysis of Covariance: age and gender) with between group post-hoc tests using multiple comparisons correction (Bonferroni) in a general linear model within the SPSS v23 framework (IBM Corporation, New York, NY, USA).

## 3. Results

AD and MCI patients demonstrated a significant increase in FDG SUVR (hyper-metabolic FDG activity) bilaterally in the cerebellum in multiple regions as shown in Fig. 1. The hyper-metabolic FDG activity in AD subjects was found in lobules VI (white arrow), IX (yellow arrow), VIII (blue arrow), and III/IV/V (cyan arrow) compared to CN participants (a). Similar regions in a FDG hyperactivity are found in AD compared to MCI subjects (b). The pattern of hyperactivity in MCI patients compared to CN subjects is similar in lobule VI and to a reduced extent in lobule III/IV/V (c). Of interest is the bilateral hyperactivity in Crus II/VIIIa (orange arrow) found in MCI patients.

**Figure 1.**
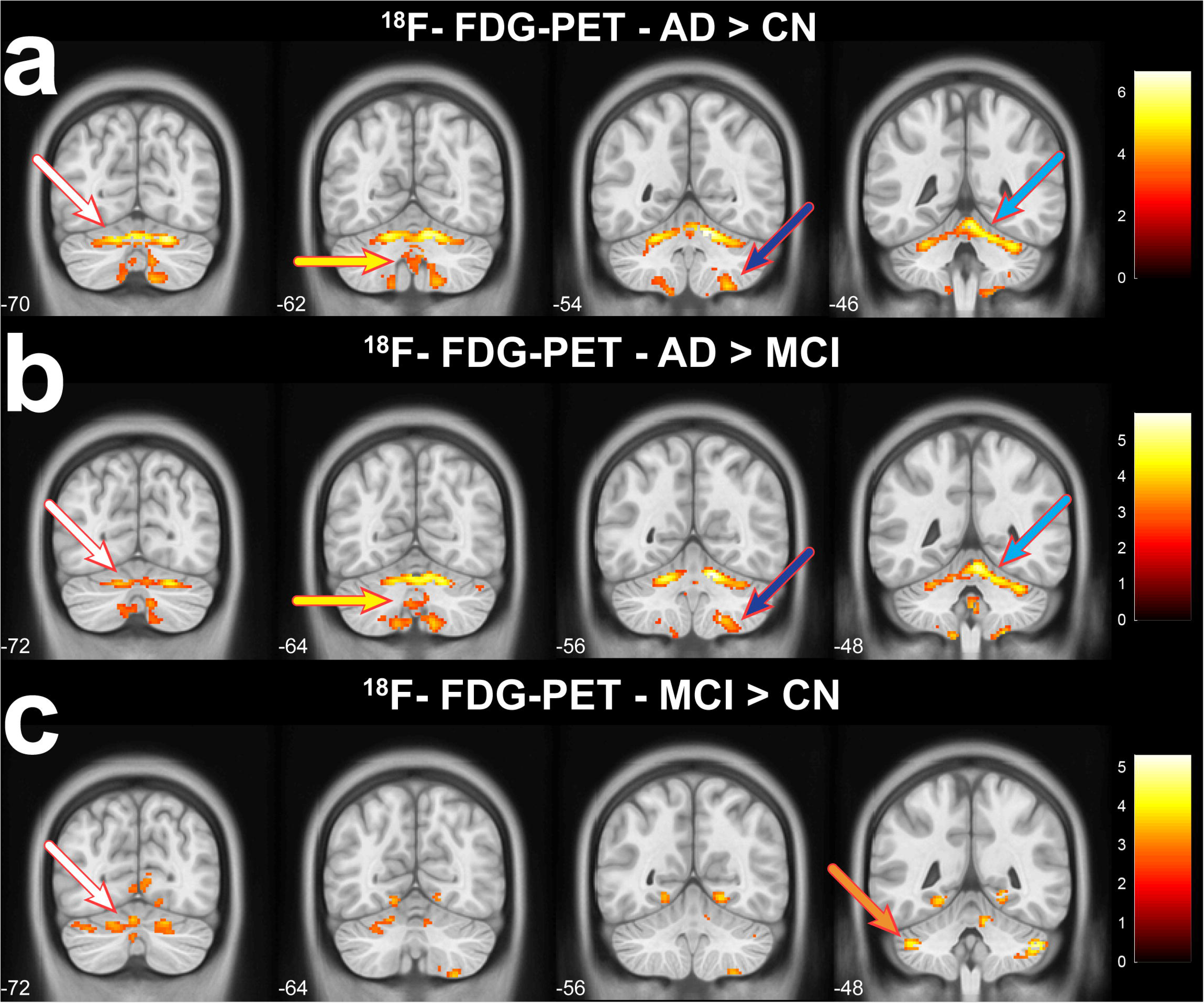
Cerebellar FDG-PET metabolic activity contrasts in the Alzheimer’s disease (N=16), mild-cognitive impaired (N=149), and cognitively normal (N=80) subjects. (a) Bilateral hyperactivity is observed in the Alzheimer’s cerebellum compared to cognitively normal patients. Hyperactivity is noted in the cerebellum lobules VI (white arrow), IX (yellow arrow), VIII (blue arrow), and III/IV/V (cyan arrow) of Alzheimer’s patients. (b) Contrast between Alzheimer’s and MCI yields similar regions of hyperactivity. (c) MCI compared to cognitively normal demonstrated a similar pattern of hyperactivity in lobule VI and to a reduced extent in lobule III/IV/V. The MCI patients have bilateral hyperactivity in Crus II/VIIIa (orange arrow). Contrasts displayed with a *p* > 0.005 and 50 voxel threshold.

As demonstrated in Fig. 2, compared to NC, hyperactivity was also found in AD in the left olfactory regions in AD, including the left lateral olfactory area (green arrow) and anterior olfactory nucleus (red arrow)(a). FDG hyperactivity is observed in the similar olfactory region in the left lateral olfactory area and anterior olfactory nucleus in MCI compared to NC participants (b). Additionally, MCI patients have a band of hyperactivity along the left olfactory tubercle/shell of the nucleus accumbens (yellow arrow).

**Figure 2.**
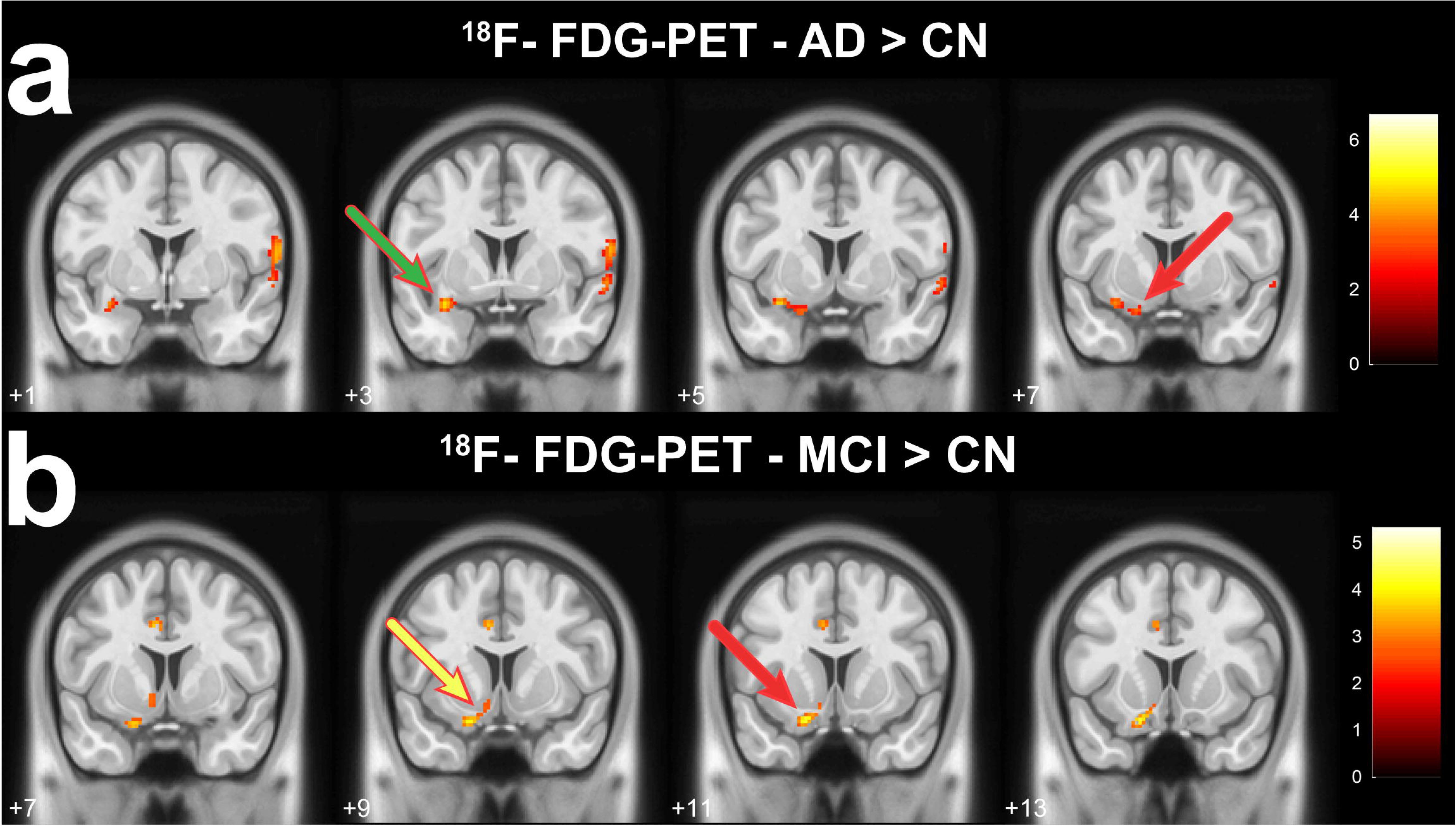
Regional mean ^18^F-FDG-SUVR contrasts in the olfactory region in Alzheimer’s disease (N=16), mild-cognitive impaired (N=149), and cognitively normal (N=80) subjects. (a) Alzheimer’s patents have FDG hyperactivity in the left lateral olfactory area (green arrow) and anterior olfactory nucleus (red arrow). (b) MCI patients have a similar pattern of olfactory region FDG hyperactivity in the left lateral olfactory area and anterior olfactory nucleus compared to cognitively normal patients. MCI patients have an additional band of hyperactivity along the left olfactory tubercle/shell of the nucleus accumbens (yellow arrow). Contrasts displayed with a *p* > 0.005 and 50 voxel threshold.

Regional FDG measures demonstrated a significantly higher FDG activity in the cerebellum of AD compared to MCI and CN participants as seen in Fig. 3. There is an increase in FDG metabolism in cerebellar lobules III, IV/V, VI, VIII, IX, and vermis of AD compared to early-, late-MCI, and cognitively normal patients. A trend of incremental FDG hypometabolism is evident in late-MCI into AD dementia. A difference between early and late-MCI patients is noted in lobule VI, with trends towards significance in other lobules. Regional measures of the primary olfactory cortex also show a trend hyper-metabolism from MCI to AD (*p* > 0.06) (Fig. 3g). Consistent with previous literature, our analysis also showed hypo-metabolism in angular gyrus in AD and MCI compared to NC subjects (Fig. 3h), as well as between AD and MCI subjects [9]. It is notable that there is not a discernable regional difference in FDG metabolism between early- and late-MCI patients, as with the cerebellar regions.

**Figure 3.**
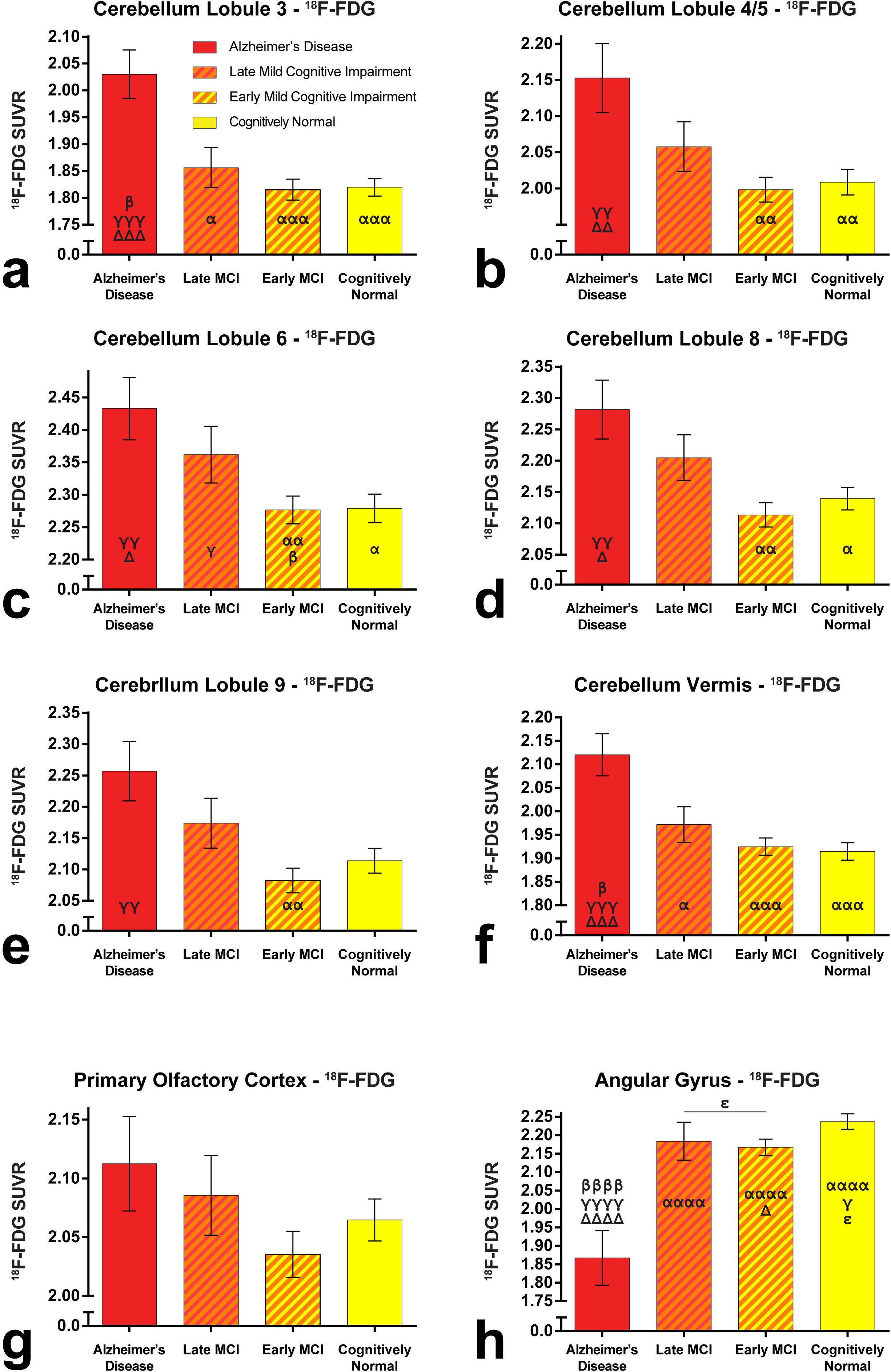
Regional brain FDG-PET metabolism in Alzheimer’s (N=16), late-MCI (N=34), early-MCI (N=115), and cognitively normal (N=80) patients. A regional increase in FDG SUVR is noted in the cerebellum of Alzheimer’s patients, congruent with the voxel-wise analysis in figure 1. Further stratifying the MCI cohort into late- and early-MCI demonstrates a linear gradated pattern in FDG hyperactivity in Lobules 4/5, VI, VIII, and IX. The primary olfactory cortex demonstrates a similar gradated regional FDG SUVE increase in early-, late-MCI, and Alzheimer’s patients, however is non-significant at *p* = 0.06. Regional FDG SUVR in the angular gyrus (H), a region of the default mode network, demonstrates an expected decrease in Alzheimer’s FDG SUVR with a gradation between AD, MCI, and CN subjects. α= difference to AD, β= difference to late-MCI, γ= difference to early-MCI, Δ= difference to cognitively normal, and ∊= difference to combined MCI. The number of indicators equates to significance level; # = *p* < 0.05, ## = *p* < 0.01, ### = *p* < 0.001, and #### = *p* < 0.0001.

FDG metabolic activity presents a complicated hypo- and hyper-activity throughout the brain. To provide a comprehensive regional FDG-SUVR measures in the brain, Fig. 4 shows 120 gray matter regions arranged radially by cortical location in AD, MCI, and CN (Fig. 4a) and early- vs. late-MCI subjects (Fig.4b). Regions with increased FDG-SUVR in AD compared to MCI and CN are clearly outlined in deep brain and cerebellar structures as well as the olfactory region in the frontal cortex. Regions with hypo-metabolism are noted in the parietal, temporal, and cingulate cortices (dashed line). Averaged MCI and CN subjects demonstrated a relative conservation of FDG-SUVR with some difference in regions aforementioned in the AD to CN comparison. Separation in FDG values in noted between the MCI group when considering early- vs. late-MCI status. There is a global trend in FDG hyper-metabolism in late-MCI patients throughout the brain with regions of corrected significance in the occipital cortex and cerebellum (Fig. 4b).

**Figure 4.**
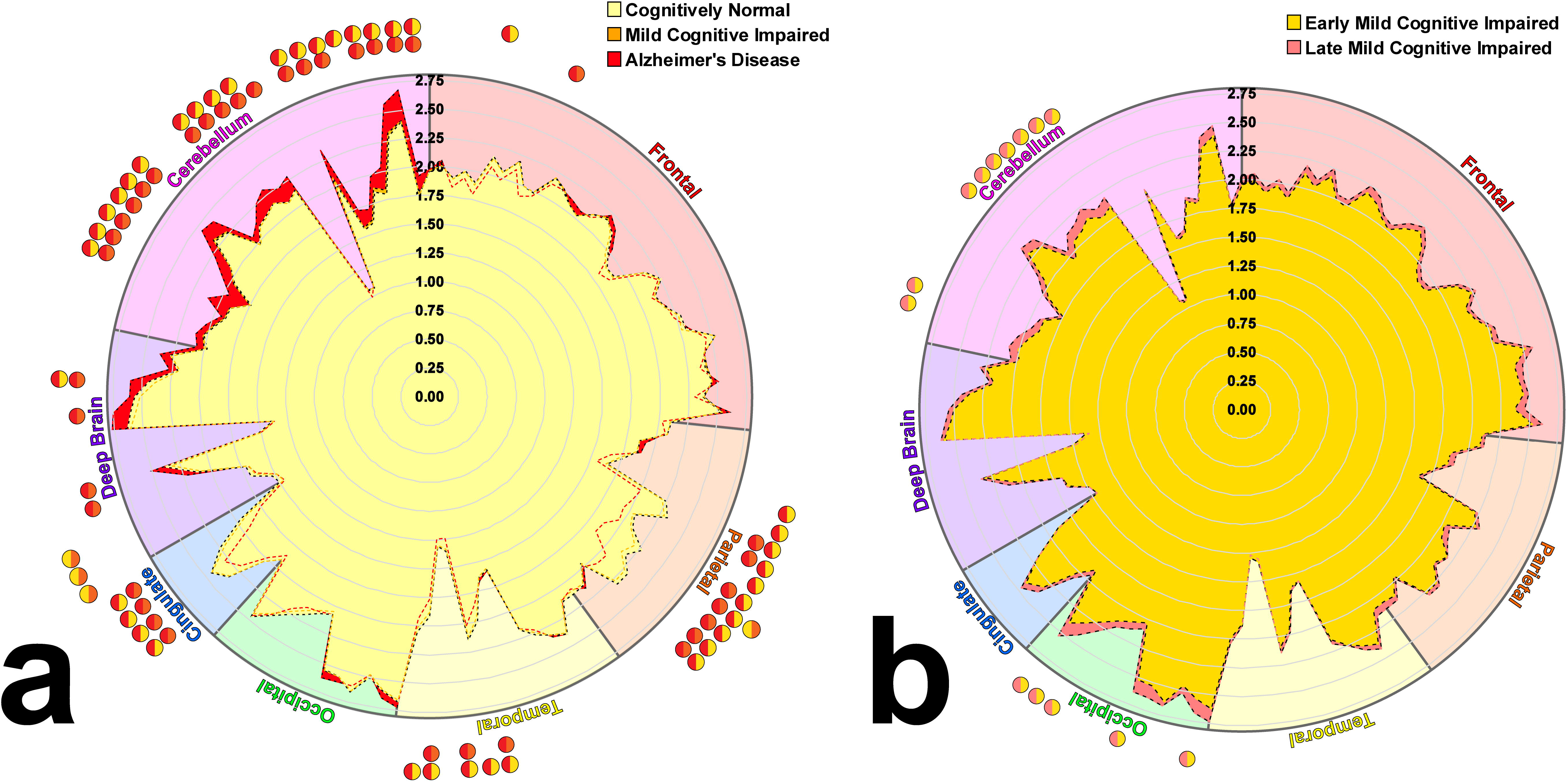
Splatterplot of FDG-PET SUVR for all 120 gray matter regions in the brain arranged radially by cortical location. Regions with significant differences (*p* < 0.05, Bonferroni MCC) are highlighted by the bi-colored circle on the outside edge indicating between group significance. Multiple regions with an increase in AD FDG metabolism are noted in the cerebellum (a). An increase in the olfactory region metabolism is noted in the frontal cortical regions. Regions with AD FDG hypo-metabolism are found the in the parietal and cingulate, indicative of the default mode regions. Further stratification of the MCI to early- and late-MCI demonstrates a global increase in FDG SUVR in the late-MCI patients, with significance in the cerebellum and occipital cortices (b).

An evaluation was conducted to compare FDG-SUVR defined using the subcortical white matter, whole cerebellum, and pons as reference regions (Fig. 5). Consistent with previously published data, Alzheimer’s hypo-FDG-metabolism was observed in the posterior cingulate, inferior parietal lobule, angular, and superior temporal gyrus regions compared to controls; all regions of the default mode network. Similar hypo-metabolic trends of FDG SUVR in the AD DMN regions are noted in the whole cerebellum and pons reference SUVR contrasts compared to the white matter as a SUVR reference.

**Figure 5.**
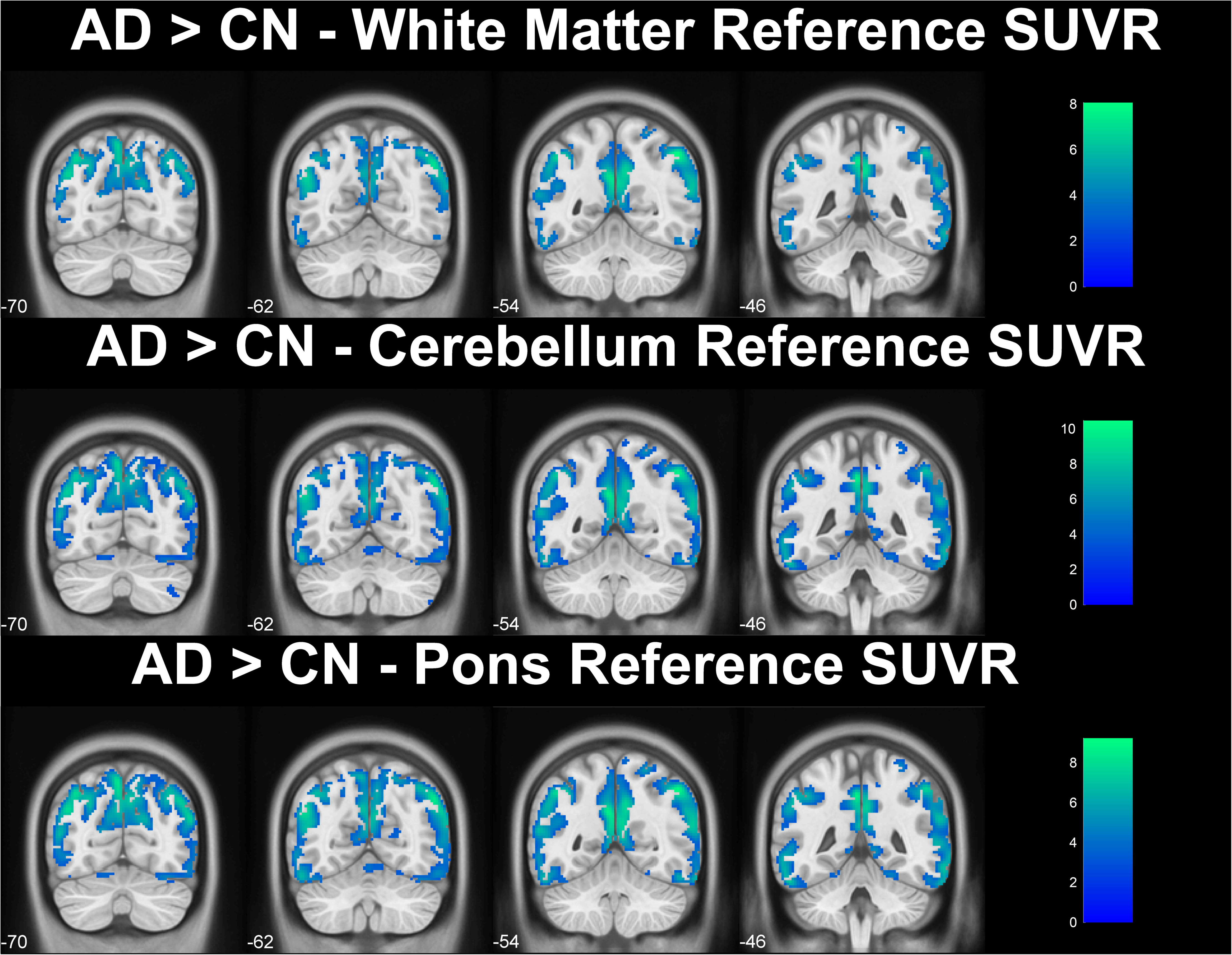
Default mode network hypo-activity in Alzheimer’s disease compared to cognitively normal patients using subcortical white matter, pons, and whole cerebellum as reference regions to calculate FDG-SUVR. The decrease in FDG metabolism is noted in regions of the default mode network, including the posterior cingulate cortex, precuneus, lateral temporal cortex, temporoparietal junction, and angular gyrus. A similar pattern of FGD hypo-metabolism is found across all three analysis methods. All contrasts displayed with a *p* < 0.001 and 50 voxel extent threshold.

## 4. Discussion

The data presented in this work revealed a regional increase in FDG metabolism in the AD cerebellum and olfactory cortical structures. To our knowledge, this is the first report demonstrating hyper-metabolism in regions of cerebellum, POC, olfactory tubercle, and nucleus accumbens in the AD brain using the same cohort. This result is intriguing as it raised a challenging question as to why these specific brain areas in AD present with hyper-metabolism while other brain regions present with hypo-metabolism due to the AD neurodegeneration.

While it has previously been straightforward to link hypo-metabolism to AD neurodegeneration. In fact, hypo FDG PET activity in these brain regions (specific regions) has been considered as a neurodegeneration marker of AD as it is correlated/hypothesized? to reduction of processes of neurons (ref). It seemed counterintuitive, however, that hyper-metabolism could also occur in AD due to neurodegeneration, especially in the cerebellum which has largely been viewed as preserved of AD pathology. Relative hyper-metabolism has been reported in a handful of neurodegenerative diseases including Parkinson’s disease [36], Amyotrophic Lateral Sclerosis [37], and MCI [38]. There is report of increased activation in anterior cerebellar regions in MCI patients compared to controls [15]. The reported hyper-metabolism from FDG measurements have been largely attributed to an increase in neuronal activity, inflammatory response, or the presence of cancerous tissue. Increased neuronal activity is also associated with behavior based focal functional activity, epileptic foci, and has been hypothesized to relate to compensatory neural activation [38]. These possible mechanisms, however, do not fit the findings here in MCI and AD.

Compared to the cerebral cortices, the cerebellum is a tri-laminar structure that has increased in size during evolutionary hominid development, paralleling that of the prefrontal and association cortices during the same phylogenetic period [18]. Importantly, AD patients were found to have an increased glucose metabolism in the cerebellum dorsally along lobules IV – VIIa, medially along lobules I – IV, and ventrally along lobules VIII- IX. The cerebellum is organized into functional networks that topographically map to association regions of the cerebrum [16]. The regions with AD hypo-metabolism correspond to functional maps of the salience, somatomotor, and DMN [16]. The cerebellar cortex largely acts through inhibitory interneuronal synapses (Purkinje, stellate, basket and Golgi cells) within the cerebellum, deep cerebellar nuclei, and projects only inhibitory afferents to the cerebrum. The local inhibitory synapses of basket cells on Purkinje cells act to disinhibit their response on deep cerebellar nuclei, upregulating their synaptic response [39]. Hyper-metabolism due to disinhibition has been noted in the cerebellum in the context of motor control [40]. Cerebellar hyper-metabolism has been observed in Parkinson’s disease in relation to cognitive impairment [36]. Hyper-metabolism in neurodegenerative disease remains a subject of further investigation.

The olfactory cortex is an evolutionary conserved paleocortex, tri-laminar in structure, and shares characteristics with lesser species three-layered cortices (i.e. reptilian) [41]. Like the cerebellum, the olfactory cortex and bulb are strongly innervated with feedforward and recurrent inhibitory circuits [42], with direct inhibitory connections projecting from the anterior cingulate cortex of the DMN to the POC [43]. These inhibitory connections are hypothesized to have a dis-inhibitory effect of excitatory neurons in the POC with distal- and inter-neuronal inhibitory connections regulating olfactory processing and perception [44]. Furthermore, local inhibitory circuits in the olfactory region are impaired by oligomeric Aβ_42_ peptide which disrupts olfactory information output and impairs GABAergic synaptic transmission in the olfactory system [45], resulting in neuronal hyperactivity [46, 47]. There has been speculation that AD pathology might initiate and spread from the entorhinal olfactory region [48, 49]. In addition, there is a known distal-inhibitory connection between the anterior cingulate cortex on the POC [43]. Taking together, the increase in glucose metabolism observed in the POC could be a result of decreased local inhibitory control of the POC coupled with a known increase in Aβ production in the entorhinal and primary olfactory cortices [50] and /or system-wised distal-inhibitory control from DMN. This finding is consistent with the proposed mechanism of hyper-metabolism in the cerebellum.

This hypothetical mechanism could explain the hypo-activity findings in the other brain areas. The nucleus accumbens and the olfactory tubercle collectively form the ventral striatum. The olfactory tubercle has three layers, similar to the rest of the olfactory system, and functionally plays a role in multisensory integration of olfactory information with other senses and reward pathways [51]. The tubercle contains GABAergic medium spiny neurons which receive glutamatergic inputs form cortical regions and dopaminergic inputs from the ventral tegmental area [52]. Neurons in the nucleus accumbens are mostly medium spiny neurons (MSNs) containing D1-excitatory, D2-inhibitory type, and GABA receptors with inhibitory control on behavior influenced by dopamine [51]. The regions also contain GABAergic interneurons which participate in powerful feedforward regional inhibition [51]. A preclinical FDG study also demonstrated an increase in FDG metabolism in the cerebellum and olfactory structures in an Alzheimer’s murine model [53]. Thus, consistent with the hypo-mentalism in the DMN, the hyper-metabolism in the olfactory structures could reflect the disruption of local and system-wised inhibitory networks due neurodegeneration of AD pathology. Unlike to the hypo-metabolism, however, the hyper-activity and metabolism in the resting-state in POC could further result in excitotoxicity and neuronal death, which, in turn, could lead to an accelerated vicious cycle of neurodegeneration. Inhibitory circuit dysfunction plays a role in the underlying cellular mechanisms underpinning AD. The impairment effect oligomeric amyloid has on inhibitory neurons has been attributed to upregulated sodium leaks with downstream deceases in action potential amplitude and frequency [54]. Inhibitory neuronal dysfunction results in large abnormal field potentials and excitability of neurons projected upon [55].

There are reports that an increase of FDG metabolism is ostensibly related to artifacts of image preprocessing in regards to selection of reference region for signal normalization [56]. Using white matter as a PET SUVR reference region has been reported to be more robust and less sensitive to age [33] and disease altering atrophy [32, 57]. Some have reported an apparent hyper-metabolism when using white matter as a reference region, with increased metabolism in the thalamus and a much more limited area of the cortex compared to whole brain normalization [58]. White matter glucose hypo-metabolism is not observed in AD [59]. An analysis of white matter FDG uptake as a function of age indicated a stable pattern of white matter glucose metabolism across the AD, MCI, or CN cohorts, concurrent with other work [33] (not shown). Furthermore, anterior and posterior cerebellar gray matter atrophy occurs during the early stages of AD, are predictors of symptom severity, and parallel cerebellar histopathological amyloid-β staging [60, 61]; potentially limiting use of the cerebellum as a normalization reference region. A comparison of the FDG SUVR maps using either whole-brain subcortical white matter, pons, or whole cerebellum as reference (Fig. 5) demonstrates that using whole-brain white matter to calculate the normalized CMRgl [32, 57] is similar in sensitivity to the other reference regions. Furthermore, the data demonstrate that subcortical white matter referencing is more sensitive than whole cerebellum referencing in detection of altered glucose metabolism in both POC and cerebellum. As it has been demonstrated previously and by our present data that the cerebellum is involved AD neurodegeneration, using it as a reference for FDG SUVR could inevitably mask the metabolic alterations due to AD. Thus, it is more advantageous to use whole-brain subcortical white matter as reference as it also minimizes the global atrophy effect due to aging [33]. The involvement of cerebellum in AD neurodegeneration [60, 61] suggests that selection of the whole-brain subcortical white matter could be a method of choice for PET SUVR for AD research.

This work demonstrates an increase in glucose utilization within the cerebellum, POC, and the ventral striatum in AD. An interpretation of observed hyper-metabolism is provided based on the inhibitory features of the olfactory system and cerebellum. The proposed mechanism, although plausible, needs to be framed in context of study limitations. The absence of balanced imaging and behavioral olfactory data within the ADNI study design does not allow correlation between observed POC hyper-metabolism and indices of olfactory function. Numerous studies in AD patients demonstrate a consistent loss of olfactory function early in the disease process and persistent throughout disease course. Thus, the significance of our data is that it generates an interesting hypothesis for further investigation. The distal and interneuronal inhibitory control of the cerebellar and olfactory systems are key factors in the cellular mechanisms that constitute the progression of AD. The hyper-metabolism in the olfactory structures and cerebellum may reflect the disruption of local and system-wide inhibitory networks due to neurodegeneration, which suggests a plausible hypothetical mechanism for the susceptibility of the olfactory system to early AD pathology. Future work focusing on inhibitory control within the olfactory structures as potential pathways of AD pathogenesis are warranted.

## Abbreviations

FDG: ^18^F-fludeoxyglucose
AFNI: Analysis of Functional NeuroImages
AD: Alzheimer’s disease
ADNI: Alzheimer’s Disease Neuroimaging Initiative
Aβ: Amyloid-beta
CN: Cognitively Normal
DMN: Default mode network
FDG-PET: Fludeoxyglucose positron emission tomography
FTD: Frontal temporal dementia
MTL: Medial temporal lobes
MCI: Mild cognitive impairment
MNI: Montreal Neurological Institute
PET: Positron emission tomography
PCC: Posterior cingulate cortex
POC: Primary Olfactory Cortex
CMRgl: Regional cerebral glucose metabolic rate
SUVR: Standardized uptake value ratio
SPM: Statistical parametric mapping

## Acknowledgements

Funding for this project has been made available in part through the **NIH (RO1-EB00454 and R01-AG027771-01A2**), the Neuroimaging Research Grant, and the Pennsylvania Department of Health using Tobacco Settlement Funds.

The authors would like to thank the patients, their families, and the ADNI researchers who graciously dedicated their time and effort to make this line of research possible.

Data collection and sharing for this project was funded by the Alzheimer’s Disease Neuroimaging Initiative (ADNI) (National Institutes of Health Grant U01 AG024904) and DOD ADNI (Department of Defense award number W81XWH-12-2-0012). ADNI is funded by the National Institute on Aging, the National Institute of Biomedical Imaging and Bioengineering, and through generous contributions from the following: AbbVie, Alzheimer’s Association; Alzheimer’s Drug Discovery Foundation; Araclon Biotech; BioClinica, Inc.; Biogen; Bristol-Myers Squibb Company; CereSpir, Inc.; Cogstate; Eisai Inc.; Elan Pharmaceuticals, Inc.; Eli Lilly and Company; EuroImmun; F. Hoffmann-La Roche Ltd and its affiliated company Genentech, Inc.; Fujirebio; GE Healthcare; IXICO Ltd.; Janssen Alzheimer Immunotherapy Research & Development, LLC.; Johnson & Johnson Pharmaceutical Research & Development LLC.; Lumosity; Lundbeck; Merck & Co., Inc.; Meso Scale Diagnostics, LLC.; NeuroRx Research; Neurotrack Technologies; Novartis Pharmaceuticals Corporation; Pfizer Inc.; Piramal Imaging; Servier; Takeda Pharmaceutical Company; and Transition Therapeutics. The Canadian Institutes of Health Research is providing funds to support ADNI clinical sites in Canada. Private sector contributions are facilitated by the Foundation for the National Institutes of Health (www.fnih.org). The grantee organization is the Northern California Institute for Research and Education, and the study is coordinated by the Alzheimer’s Therapeutic Research Institute at the University of Southern California. ADNI data are disseminated by the Laboratory for Neuro Imaging at the University of Southern California.

## References

[1] Jack CR, Jr., Bennett DA, Blennow K, Carrillo MC, Dunn B, Haeberlein SB, et al. NIA-AA Research Framework: Toward a biological definition of Alzheimer’s disease. Alzheimers Dement. 2018;14:535–62.

[2] Lehmann M, Ghosh PM, Madison C, Laforce R, Jr., Corbetta-Rastelli C, Weiner MW, et al. Diverging patterns of amyloid deposition and hypometabolism in clinical variants of probable Alzheimer’s disease. Brain. 2013;136:844–58.

[3] Foster NL, Heidebrink JL, Clark CM, Jagust WJ, Arnold SE, Barbas NR, et al. FDG-PET improves accuracy in distinguishing frontotemporal dementia and Alzheimer’s disease. Brain. 2007;130:2616–35.

[4] Mosconi L. Brain glucose metabolism in the early and specific diagnosis of Alzheimer’s disease. FDG-PET studies in MCI and AD. Eur J Nucl Med Mol Imaging. 2005;32:486–510.

[5] Mosconi L, De Santi S, Li Y, Li J, Zhan J, Tsui WH, et al. Visual rating of medial temporal lobe metabolism in mild cognitive impairment and Alzheimer’s disease using FDG-PET. Eur J Nucl Med Mol Imaging. 2006;33:210–21.

[6] Bokde AL, Pietrini P, Ibanez V, Furey ML, Alexander GE, Graff-Radford NR, et al. The effect of brain atrophy on cerebral hypometabolism in the visual variant of Alzheimer disease. Arch Neurol. 2001;58:480–6.

[7] Mosconi L, Sorbi S, de Leon MJ, Li Y, Nacmias B, Myoung PS, et al. Hypometabolism exceeds atrophy in presymptomatic early-onset familial Alzheimer’s disease. J Nucl Med. 2006;47:1778–86.

[8] Rodriguez-Oroz MC, Gago B, Clavero P, Delgado-Alvarado M, Garcia-Garcia D, Jimenez-Urbieta H. The relationship between atrophy and hypometabolism: is it regionally dependent in dementias? Curr Neurol Neurosci Rep. 2015;15:44.

[9] Marcus C, Mena E, Subramaniam RM. Brain PET in the diagnosis of Alzheimer’s disease. Clin Nucl Med. 2014;39:e413–22; quiz e23-6.

[10] Greicius MD, Srivastava G, Reiss AL, Menon V. Default-mode network activity distinguishes Alzheimer’s disease from healthy aging: evidence from functional MRI. Proc Natl Acad Sci U S A. 2004;101:4637–42.

[11] Marchitelli R, Aiello M, Cachia A, Quarantelli M, Cavaliere C, Postiglione A, et al. Simultaneous resting-state FDG-PET/fMRI in Alzheimer Disease: Relationship between glucose metabolism and intrinsic activity. Neuroimage. 2018;176:246–58.

[12] Sjobeck M, Englund E. Alzheimer’s disease and the cerebellum: a morphologic study on neuronal and glial changes. Dement Geriatr Cogn Disord. 2001;12:211–8.

[13] Mann DM, Jones D, Prinja D, Purkiss MS. The prevalence of amyloid (A4) protein deposits within the cerebral and cerebellar cortex in Down’s syndrome and Alzheimer’s disease. Acta Neuropathol. 1990;80:318–27.

[14] Guo CC, Tan R, Hodges JR, Hu X, Sami S, Hornberger M. Network-selective vulnerability of the human cerebellum to Alzheimer’s disease and frontotemporal dementia. Brain. 2016;139:1527–38.

[15] Jacobs HIL, Hopkins DA, Mayrhofer HC, Bruner E, van Leeuwen FW, Raaijmakers W, et al. The cerebellum in Alzheimer’s disease: evaluating its role in cognitive decline. Brain. 2018;141:37–47.

[16] Buckner RL, Krienen FM, Castellanos A, Diaz JC, Yeo BT. The organization of the human cerebellum estimated by intrinsic functional connectivity. J Neurophysiol. 2011;106:2322–45.

[17] Zheng W, Liu X, Song H, Li K, Wang Z. Altered Functional Connectivity of Cognitive-Related Cerebellar Subregions in Alzheimer’s Disease. Front Aging Neurosci. 2017;9:143.

[18] Klein AP, Ulmer JL, Quinet SA, Mathews V, Mark LP. Nonmotor Functions of the Cerebellum: An Introduction. AJNR Am J Neuroradiol. 2016;37:1005–9.

[19] Doty RL. Olfactory deficit in Alzheimer’s disease? Am J Psychiatry. 2001;158:1533–4; author reply 4-5.

[20] Martinez B, Karunanayaka P, Wang J, Tobia MJ, Vasavada M, Eslinger PJ, et al. Different patterns of age-related central olfactory decline in men and women as quantified by olfactory fMRI. Oncotarget. 2017;8:79212–22.

[21] Wang J, Eslinger PJ, Doty RL, Zimmerman EK, Grunfeld R, Sun X, et al. Olfactory deficit detected by fMRI in early Alzheimer’s disease. Brain Res. 2010;1357:184–94.

[22] Serby M, Mohan C, Aryan M, Williams L, Mohs RC, Davis KL. Olfactory identification deficits in relatives of Alzheimer’s disease patients. Biol Psychiatry. 1996;39:375–7.

[23] Oleson S, Murphy C. Olfactory Dysfunction in ApoE varepsilon4/4 Homozygotes with Alzheimer’s Disease. J Alzheimers Dis. 2015;46:791–803.

[24] Roberts RO, Christianson TJ, Kremers WK, Mielke MM, Machulda MM, Vassilaki M, et al. Association Between Olfactory Dysfunction and Amnestic Mild Cognitive Impairment and Alzheimer Disease Dementia. JAMA Neurol. 2016;73:93–101.

[25] Braak H, Alafuzoff I, Arzberger T, Kretzschmar H, Del Tredici K. Staging of Alzheimer disease-associated neurofibrillary pathology using paraffin sections and immunocytochemistry. Acta Neuropathol (Berl). 2006.

[26] Vasavada M, Zhang H, Wang J, Karunanayaka P, Eslinger PJ, Gill DJ, et al. Functional connectivity of the primary olfactory cortex is decreased in Alzheimer’s disease and mild cognitive impairment. Proceedings 22nd Scientific Meeting, International Society for Magnetic Resonance in Medicine. Milan2014. p. 0490.

[27] Karunanayaka PR, Wilson DA, Tobia MJ, Martinez BE, Meadowcroft MD, Eslinger PJ, et al. Default mode network deactivation during odor-visual association. Hum Brain Mapp. 2017;38:1125–39.

[28] Alzheimer’s Disease Neuroimaging Initiative – Study Documents. http://adni.loni.usc.edu/methods/documents/.

[29] Weiner MW. ADNI Overview. http://citeseerxistpsuedu/viewdoc/download?doi=10116572042&rep=rep1&type=pdf.

[30] Cox RW. AFNI: what a long strange trip it’s been. Neuroimage. 2012;62:743–7.

[31] Thomas BA, Cuplov V, Bousse A, Mendes A, Thielemans K, Hutton BF, et al. PETPVC: a toolbox for performing partial volume correction techniques in positron emission tomography. Phys Med Biol. 2016;61:7975–93.

[32] Blautzik J, Brendel M, Sauerbeck J, Kotz S, Scheiwein F, Bartenstein P, et al. Reference region selection and the association between the rate of amyloid accumulation over time and the baseline amyloid burden. Eur J Nucl Med Mol Imaging. 2017;44:1364–74.

[33] Borghammer P, Jonsdottir KY, Cumming P, Ostergaard K, Vang K, Ashkanian M, et al. Normalization in PET group comparison studies--the importance of a valid reference region. Neuroimage. 2008;40:529–40.

[34] Brett M, Anton J-L, Valabregue R, Poline JB. Region of interest analysis using an SPM toolbox. 8th International Conference on Functional Mapping of the Human Brain. Sendai, Japan2002.

[35] Rolls ET, Joliot M, Tzourio-Mazoyer N. Implementation of a new parcellation of the orbitofrontal cortex in the automated anatomical labeling atlas. Neuroimage. 2015;122:1–5.

[36] Blum D, la Fougere C, Pilotto A, Maetzler W, Berg D, Reimold M, et al. Hypermetabolism in the cerebellum and brainstem and cortical hypometabolism are independently associated with cognitive impairment in Parkinson’s disease. Eur J Nucl Med Mol Imaging. 2018.

[37] Cistaro A, Valentini MC, Chio A, Nobili F, Calvo A, Moglia C, et al. Brain hypermetabolism in amyotrophic lateral sclerosis: a FDG PET study in ALS of spinal and bulbar onset. Eur J Nucl Med Mol Imaging. 2012;39:251–9.

[38] Ashraf A, Fan Z, Brooks DJ, Edison P. Cortical hypermetabolism in MCI subjects: a compensatory mechanism? Eur J Nucl Med Mol Imaging. 2015;42:447–58.

[39] Purves D, Augustine GJ, Fitzpatrick D. Neuroscience. Sixth edition. ed: Oxford University Press; 2017.

[40] Mink JW. The basal ganglia: focused selection and inhibition of competing motor programs. Prog Neurobiol. 1996;50:381–425.

[41] Klingler E. Development and Organization of the Evolutionarily Conserved Three-Layered Olfactory Cortex. eNeuro. 2017;4.

[42] Large AM, Vogler NW, Mielo S, Oswald AM. Balanced feedforward inhibition and dominant recurrent inhibition in olfactory cortex. Proc Natl Acad Sci U S A. 2016;113:2276–81.

[43] Garcia-Cabezas MA, Barbas H. A direct anterior cingulate pathway to the primate primary olfactory cortex may control attention to olfaction. Brain Struct Funct. 2014;219:1735–54.

[44] Cao LH, Yang D, Wu W, Zeng X, Jing BY, Li MT, et al. Odor-evoked inhibition of olfactory sensory neurons drives olfactory perception in Drosophila. Nat Commun. 2017;8:1357.

[45] Hu B, Geng C, Hou XY. Oligomeric amyloid-beta peptide disrupts olfactory information output by impairment of local inhibitory circuits in rat olfactory bulb. Neurobiol Aging. 2017;51:113–21.

[46] Busche MA, Eichhoff G, Adelsberger H, Abramowski D, Wiederhold KH, Haass C, et al. Clusters of hyperactive neurons near amyloid plaques in a mouse model of Alzheimer’s disease. Science. 2008;321:1686–9.

[47] Busche MA, Konnerth A. Neuronal hyperactivity--A key defect in Alzheimer’s disease? Bioessays. 2015;37:624–32.

[48] Franks KH, Chuah MI, King AE, Vickers JC. Connectivity of Pathology: The Olfactory System as a Model for Network-Driven Mechanisms of Alzheimer’s Disease Pathogenesis. Front Aging Neurosci. 2015;7:234.

[49] Liu X, Zeng K, Li M, Wang Q, Liu R, Zhang B, et al. Expression of P301L-hTau in mouse MEC induces hippocampus-dependent memory deficit. Sci Rep. 2017;7:3914.

[50] Wu N, Rao X, Gao Y, Wang J, Xu F. Amyloid-beta deposition and olfactory dysfunction in an Alzheimer’s disease model. J Alzheimers Dis. 2013;37:699–712.

[51] Wesson DW, Wilson DA. Sniffing out the contributions of the olfactory tubercle to the sense of smell: hedonics, sensory integration, and more? Neurosci Biobehav Rev. 2011;35:655–68.

[52] Ikemoto S. Brain reward circuitry beyond the mesolimbic dopamine system: a neurobiological theory. Neurosci Biobehav Rev. 2010;35:129–50.

[53] Lu Y, Ren J, Cui S, Chen J, Huang Y, Tang C, et al. Cerebral Glucose Metabolism Assessment in Rat Models of Alzheimer’s Disease: An 18F-FDG-PET Study. Am J Alzheimers Dis Other Demen. 2016;31:333–40.

[54] Perez C, Ziburkus J, Ullah G. Analyzing and Modeling the Dysfunction of Inhibitory Neurons in Alzheimer’s Disease. PLoS One. 2016;11:e0168800.

[55] Hazra A, Gu F, Aulakh A, Berridge C, Eriksen JL, Ziburkus J. Inhibitory neuron and hippocampal circuit dysfunction in an aged mouse model of Alzheimer’s disease. PLoS One. 2013;8:e64318.

[56] Borghammer P, Cumming P, Aanerud J, Gjedde A. Artefactual subcortical hyperperfusion in PET studies normalized to global mean: lessons from Parkinson’s disease. Neuroimage. 2009;45:249–57.

[57] Rasmussen JM, Lakatos A, van Erp TG, Kruggel F, Keator DB, Fallon JT, et al. Empirical derivation of the reference region for computing diagnostic sensitive (1)(8)fluorodeoxyglucose ratios in Alzheimer’s disease based on the ADNI sample. Biochim Biophys Acta. 2012;1822:457–66.

[58] Lee SJ, Lee WY, Kim YK, An YS, Cho JW, Choi JY, et al. Apparent relative hypermetabolism of selective brain areas in Huntington disease and importance of reference region for analysis. Clin Nucl Med. 2012;37:663–8.

[59] Jeong YJ, Yoon HJ, Kang DY. Assessment of change in glucose metabolism in white matter of amyloid-positive patients with Alzheimer disease using F-18 FDG PET. Medicine (Baltimore). 2017;96:e9042.

[60] Bruchhage M, Correla S, Malloy P, Salloway S, Deoni S. Using machine learning to classify early stages of cognitive decline from typical ageing – the cerebellum more than just a bystander. Proc Intl Soc Mag Reson Med. Paris, France2018.

[61] Thal DR, Rub U, Orantes M, Braak H. Phases of A beta-deposition in the human brain and its relevance for the development of AD. Neurology. 2002;58:1791–800.

